# Divergent transcriptional response to thermal stress among life stages could constrain coral adaptation to climate change

**DOI:** 10.1101/2022.04.29.490056

**Authors:** Maria Ruggeri, Yingqi Zhang, Galina V. Aglyamova, Carly D. Kenkel

## Abstract

The ability for adaptation to keep pace with environmental change largely depends on how efficiently selection can act on heritable genetic variation. Complex life cycles may either promote or constrain adaptation depending on the integration or independence of fitness-related traits over development. Reef-building corals exhibit complex life cycles and are sensitive to increasing temperatures, highlighting the need to understand the heritable potential of the thermal stress response and how it is regulated over development. Here we used tag-based RNA-seq to profile global gene expression in inshore and offshore *P. astreoides* adults and their offspring recruits in response to a 16-day heat stress, and larvae from separate families in response to a 4-day heat stress, to test whether gene expression patterns differentiating adult populations, and potentially underlying differences in thermal tolerance, persist in thermally naive life stages. Host developmental stage had a major effect on both host and symbiont expression, despite symbionts being directly inherited from parent colonies, and modulated the response to thermal stress, suggesting the holobiont response to selection varies across life stages. Populations also exhibited origin-specific treatment responses, but the magnitude of the response differed among populations and life stages. Inshore parents and their juvenile offspring exhibited a more robust response to heat stress compared to offshore-origin corals, indicating expression plasticity may be heritable. However, larval populations exhibited the opposite response, possibly due to stage-specific differences or exposure duration. Overall, this study shows that putatively adaptive regulatory variation can be heritable, but the identity of thermally responsive genes are stage-specific, which will have major implications for predicting the evolutionary response of corals in a changing environment.

## Introduction

Central to predicting organismal responses to climate change is understanding how fitness landscapes will change over time. If traits are heritable and unconstrained, adaptation may be able to keep pace with environmental change (Bay et al., 2017a). However, organisms with complex life cycles can occupy different forms, functions, and even habitats within a single lifetime (Wilbur, 1980). As life stages share a common genome, selection pressures that vary over the course of development (Aguirre et al., 2014; Coronado-Zamora et al., 2019a) can affect total organismal fitness and, ultimately, adaptation (Albecker et al., 2021). Gene expression serves as a mechanistic link between genotype and phenotype and can be measured over development to explore the degree of integration or independence between discrete life stages. Differential expression is common between developmental transitions (Herrig et al., 2021; Reyes-Bermudez et al., 2016; Schmid et al., 2005), however, it is unclear whether traits relevant to all life stages, such as the molecular response to stress, also vary over development. As global temperatures continue to increase, selective pressures will also intensify in thermally vulnerable species (Bell & Collins, 2008). Understanding how the thermal stress response is modulated over development is key to predicting whether a complex life cycle will constrain or promote adaptation to the novel environments expected under climate change.

Autonomy, or ontogenetic decoupling, is one hypothesis for the evolution of discrete developmental stages, where genes can evolve independently due to stage-specific selective pressures (Moran, 1994). Alternatively, traits can be genetically correlated or expressed across multiple life stages, conferring either a fitness advantage (synergistic pleiotropy) or disadvantage (antagonistic pleiotropy) in subsequent stages (Conner, 2012; Schluter et al., 1991). Synergistic pleiotropy results in consistent selection across all life stages and is thus predicted to result in rapid evolution (Albecker et al., 2021). Genes that are only expressed in discrete life stages, such as in ontogenetic decoupling, are predicted to adapt slower than those with synergistic pleiotropy but faster than those with antagonistic pleiotropy, where expression will differentially affect fitness over developmental time and constrain adaptation (Albecker et al., 2021)

Reef-building corals, the foundation of tropical reef ecosystems, have complex life cycles and are extremely sensitive to rising temperature. Like many marine invertebrates, coral development involves dramatic metamorphosis, where the body plan is restructured from a free-swimming larva into a sessile, juvenile polyp (Richmond & Hunter, 1990). As juveniles, coral begin to form a calcium carbonate skeleton and invest in colony growth until they reach the reproductively mature, adult stage. Corals also form a nutritive symbiosis with dinoflagellates in the family Symbiodiniaceae (Muscatine, 1990), which are either acquired directly from maternal colonies during oogenesis (vertical transmission) or from the external environment (horizontal transmission) during the larval or polyp stage. Although this partnership is mutually beneficial under normal conditions, the loss of algal cells or their photosynthetic pigments, known as coral bleaching, occurs only 1-2°C above local summer temperatures (Strong et al., 2011), and can lead to starvation and eventual death of the host under prolonged stress. Increasingly frequent and severe thermal anomalies are resulting in mass bleaching and mortality events on a worldwide scale (Eakin et al., 2019; Hoegh-Guldberg, 1999; Hughes et al., 2018). Despite the importance of early life stages in the demographic recovery from such events (Doropoulos et al., 2015; Hughes & Tanner, 2000), only 2% of all coral heat stress experiments have been performed on larvae and 1% on juvenile recruits, and even fewer studies have included more than one life stage (Rowan et al., 2020). Monitoring of coral populations over natural bleaching events suggests that adults may differ from earlier life stages in their susceptibility to thermal stress (Alvarez-Noriega et al, 2018), emphasizing the need for comparative, mechanistic studies across multiple life stages.

Earlier work examining gene regulation over the course of coral development suggests that life stages may express unique parts of the genome (Reyes-Bermudez et al., 2016; Schwarz et al., 2008). However, these prior studies focused on identifying differences during major developmental transitions, such as the onset of symbiosis (Schwarz et al., 2008), metamorphosis, and calcification (Reyes-Bermudez et al., 2016), while only one study examined the molecular response to a common stressor, salinity, across multiple life stages (Aguilar et al. 2019; Kenkel and Wright, *in press*). In a meta-analysis of expression response to stress in *Acroporid* corals, Dixon et al (2020) found life stage to be the major factor differentiating transcriptional variation, but most studies were limited to a single life stage, precluding any formal investigation of life-stage differences. However, developmental stage has been found to modulate the transcriptional response to stress in other systems, including water and salinity stress in plants (Garg et al., 2016; Liu et al., 2021), viral infection in insects (Schneweis et al., 2017), and pH stress in sea urchins (Devens et al., 2020). It is therefore necessary to explore how functional variation is regulated across different developmental stages to inform understanding of coral adaptive capacity.

Contemporary populations that are locally adapted to thermally variable environments represent promising study systems for investigating whether patterns of regulation driving thermal tolerance are heritable. Coral populations from backreef pools in American Samoa and nearshore reefs in the Florida Keys exhibit local thermal adaptation, with more thermally variable reefs having more tolerant adults (Kenkel, Goodbody-Gringley, et al., 2013; Palumbi et al., 2014), possibly driven by differences in gene expression plasticity (Barshis et al., 2013a; Kenkel & Matz, 2016). Corals from highly variable backreef pools constitutively up or down-regulate stress responsive genes in anticipation of thermal stress, known as front or backloading, and thus have reduced expression plasticity compared to more susceptible corals originating from less variable pools (Barshis et al., 2013b). In contrast, locally adapted corals from more variable, inshore reefs in the Florida Keys exhibit greater plasticity of the environmental stress response and reduced bleaching compared to offshore-origin corals (Kenkel, Goodbody-Gringley, et al., 2013; Kenkel & Matz, 2016). Despite these different regulatory responses, evidence for population-level variation in expression plasticity suggests that plasticity may be heritable and selected for across generations, leading to local thermal adaptation. However, these hypotheses have only been tested in adult corals, where the effect of long-term acclimatization cannot be teased apart from genetic adaptation. There is, however, evidence that parental colonies can modulate the molecular and physiological response to thermal stress (Dixon et al., 2015a; Kirk et al., 2018), suggesting that thermal tolerance heritability could be explained by inheritance of gene regulatory elements and differentially selected for across populations.

Here, we compare the molecular response to heat stress across three different life stages of *Porites astreoides* and their symbionts from inshore and offshore reefs in the Florida Keys to understand how the thermal stress response is regulated during development and whether parental patterns of regulation are inherited by offspring. *P. astreoides* is a brooding coral, meaning gamete fertilization occurs internally and competent planula larvae are released which then settle and metamorphose into juvenile recruits (Richmond & Hunter, 1990). All independent life stages are symbiotic, with symbionts being vertically inherited from maternal colonies (Thornhill et al., 2006). Both adults and larvae from inshore reefs were previously found to bleach less than those from offshore locations in response to thermal stress (Kenkel et al., 2015; Zhang et al., 2019). Although adult populations and their symbionts differ in the magnitude of gene expression plasticity, which could underpin differences in tolerance (Kenkel & Matz, 2016), it is unknown whether larvae and juveniles also exhibit these patterns of regulation. In this study, adult colonies from inshore and offshore populations and their recruit offspring, as well as larvae from separate families, were profiled for gene expression after exposure to controlled thermal stress experiments to ask 1) What are the effects of developmental stage and reef origin on host and symbiont expression? 2) How does the host developmental stage affect the molecular response to heat stress? and 3) Are patterns of regulation differentiating inshore and offshore adult populations present in thermally naïve life stages?

## Methods

### Thermal stress experiments

Adults and their recruit offspring were subject to a 16-day common garden heat stress experiment as described in Kenkel et al. (2015). Briefly, adult colonies were collected from one inshore and one offshore reef site in April 2012 during peak larval release and held at Mote Marine Laboratory’s Tropical Research Lab in Summerland Key, FL under permit #FKNMS-2012-028. Larvae released from parent colonies were settled onto terra cotta tiles to obtain families of juvenile recruits and parent colonies were halved using a tile saw after spawning Parent colonies and their recruits were then reared in a common garden environment for 5 weeks. After the acclimation period, paired parent colony halves and recruits were distributed into experimental tanks and exposed to either control (28 +/-0.4°C) or elevated temperatures (30.9 +/-1.1°C) for 14 days. Samples from 7 inshore and 5 offshore families were then flash frozen in liquid nitrogen and stored at −80°C until RNA extractions.

Larvae were sampled for gene expression analysis after being exposed to a moderate heat stress for 4 days as described in Zhang et al. (2019). In April 2018 parent colonies were collected from the same inshore and offshore reef sites used in the 2012 experiment and brought to Mote Marine Laboratory’s IC2R3 in Summerland Key, FL under permit #FKNMS-2018-033. Larvae were collected from each parent colony (3 inshore and 4 offshore) and separated by family in floating netwells, with 10 larvae per family per netwell across 3 replicate netwells per temperature treatment. Temperature was then ramped up to 32°C over 24 hours in the thermal stress treatment, while the control treatment remained at room temperature (24°C). Larvae were sampled after 4 days of exposure and frozen at −80°C until processing.

### RNA isolation, library preparation, and sequencing

Extraction and library preparation protocols for the adult/recruit samples follow those described in Kenkel & Matz (2016). Briefly, an RNAqueous kit (Ambion, Life Technologies) was used to extract total RNA. Samples were subsequently DNAse treated according to the RNAqueous 4-PCR kit protocol, after which the concentration of RNA was estimated using the Nanodrop 2000 (Thermo-Fisher). Total RNA was extracted from the larval samples using an Aurum Total RNA Mini Kit (Bio-Rad, USA) and DNAse treated using the on-column protocol as described in the manufacturers instructions. RNA concentration was estimated using a Take2 plate on a Synergy H1 microplate reader (Biotek). For both sample sets, one microgram of total RNA per sample was used to generate tag-based RNA-seq, or TagSeq, libraries (Lohman et al., 2016; Meyer et al., 2011), with modifications for sequencing on the Illumina platform as described at https://github.com/ckenkel/tag-based_RNAseq.

Adult/recruit and larval TagSeq libraries were sequenced on an Illumina HiSeq 2500 independently, in 2013 by the Genomic Sequencing and Analysis Facility at UT Austin and 2018 by the USC Genome Core, respectively. Adult/recruit libraries were sequenced at an average depth of 6.27 million reads per sample (median = 5.89; range = 0.23-17.4) and larval libraries were sequenced at an average depth of 5.60 million reads per sample (median 5.16; range = 3.06-10.61). Raw sequence data can be found under NCBI BioProject PRJNA666709.

### Data processing

Data cleaning and processing was performed on USC’s Center for Advanced Research Computing (CARC) following the bioinformatic protocols described in https://github.com/ckenkel/tag-based_RNAseq. First, PCR duplicates and reads missing adaptor sequences were removed using custom perl scripts. Adaptor sequences, poly-A tail residues and sample-specific barcodes were trimmed, and reads were then filtered to retain only those exhibiting a quality score of 20 (99% probability of correct base call) over 70% of the read. After cleaning and filtering, SHRiMP (Rumble et al., 2009) was used to competitively map reads to a concatenated *Porites astreoides* host and *Symbiodinium* spp. (KB8, formerly Clade A) transcriptome (Bayer et al., 2012; Mansour et al., 2016). A custom perl script was then used to sum the number of reads mapping to each isogroup by reference. The resulting count data was separated by isogroup into a host and symbiont data frame and analyzed separately.

### Differential gene expression analysis

The package DESeq2 (Love et al., 2014) in R (v3.6.1) was used to statistically evaluate differential expression across life stages, reef origin, and treatment groups. Differential expression analyses were performed separately for adult/recruit and larval datasets. Scripts and input files for DESeq2 and downstream analyses can be found at https://github.com/mruggeri55/PastGE-lifestage. Outliers were first screened using the R package arrayQualityMetrics (Kauffmann et al., 2009) and genes with low abundance transcripts (count <2 in 90% of samples) were filtered out. After filtering, 20,185 and 14,124 high-expression genes remained for adult/recruit host and symbiont datasets respectively. For the host larval expression, 23,806 genes remained. As only ∼3,000 genes remained after filtering, symbiont expression in the larval dataset was excluded from further analyses. Variance stabilizing transformation (VST) was applied to count data for principal components analysis (PCA) to visualize the dominant factors driving transcriptional variation.

DESeq2 was used to contrast gene expression based on life stage, origin, and treatment (∼ stage + origin + treatment) using the default settings. Significance testing was determined using a Wald test after independent filtering using an FDR threshold of 0.1. Multiple test correction was applied to raw p-values following (Benjamini & Hochberg, 1995) and adjusted p-values less than 0.1 were deemed significant. Specific contrasts were also used to evaluate the effects of life stage or reef origin on the treatment response. Contrasts were created by using a grouping variable of life stage and treatment (i.e. recruit-control, recruit-heat, adult-control, adult-heat) while controlling for origin effects (∼ origin + group) in order to evaluate the effects of life stage on the treatment response. A separate model was created to contrast the treatment response according to reef origin while controlling for life stage (∼ stage + group).

### Front and backloading analysis

To test whether treatment inducible genes in one population or life stage were not detected in the other due to front/backloading or differences in variance, we generated a correlation plot of the log fold change (LFC) across treatment groups for significant differentially expressed genes following (Barshis et al., 2013b). Genes deviating from the one to one line indicate potential candidates for front/backloading whereas those along the one to one line responded similarly in both groups and, therefore, were likely not detected due to differences in variance among groups. Candidate front/backloaded genes were identified as those that were significantly differentially expressed in control conditions (constitutively up/down-regulated) and did not respond significantly to treatment in one group, but were significantly differentially expressed in another group. This analysis was repeated across life stages and reef origin for both the host and symbiont.

### Discriminant analysis

Discriminant analysis of principal components was performed using the R package adegenet (Jombart, 2008). To explore shifts in expression profiles based on population, the discriminant function was defined by contrasting inshore samples in control and heat conditions including all treatment responsive genes. The number of PCs used to create the function were chosen to capture at least 80% of transcriptional variance. The function was then applied to offshore samples and plotted on the same axis as the inshore response. MCMCglmm (Hadfield, 2010) was used to model the population by treatment interaction to determine whether one population had a significantly greater response to treatment. This analysis was also repeated by contrasting all treatment responsive genes in offshore control and heated conditions and applying the function to inshore samples. In order to determine whether differences in plasticity underlie differences in bleaching phenotypes, a linear model was used to test whether the distance along the discriminant function for each family was correlated to changes in previously published bleaching phenotypes, including larval chlorophyll content (Zhang et al., 2019) and adult bleaching score (Kenkel et al., 2015).

### Functional enrichment

Gene ontology annotations were performed by blasting nucleotide sequences to the SwissProt-UniProt database (Boutet et al., 2007) allowing a maximum of 5 alignments and retaining hits with a minimum e-value of 0.0001. Rank-based gene ontology (GO) enrichments were then performed on signed log p-values for each DESEq2 contrast using the package GO_MWU (Wright et al., 2015) with a FDR threshold of 10%. Protein sequences were further categorized into EuKaryotic Orthologous Groups (KOG) using the online interface of Eggnog-mapper v5.0 (Cantalapiedra et al., 2021) with default settings. Contigs belonging to more than one KOG class were excluded from further analysis. In cases where a single isogroup was made up of multiple contigs with variable KOG annotations, the class that made up the majority of contigs was retained for that isogroup. If no class was in the majority, the annotation with the best blast score was chosen. The KOGMWU R package (Dixon et al., 2015b) was then used to calculate delta ranks of KOG classes across life stages based on their treatment response. To test whether KOG delta ranks were correlated across life stages, pairwise Pearson correlations were calculated and visualized using the R package ‘corrplot’.

## Results

### Life stage drives expression variation in both hosts and symbionts

Life history stage was the primary driver of transcriptional variation in both the coral hosts and dinoflagellate endosymbionts. Adult and recruit samples separated by life stage along the first principal component of variance stabilized counts for both the host (Fig 1A) and symbiont (Fig 1B) explaining 31% and 28% of transcriptional variance, respectively. Life stage also accounted for the highest proportion of differentially expressed genes, with 78.5% (6,944) of all host genes (8,851) differentially expressed between adults and recruits (Fig S1a). Adult corals more highly expressed genes involved in proteolysis, protein folding, immune system processes, and ossification compared to juvenile recruits (Table S1). In contrast, recruits overexpressed signal transduction, ion transport, system processes, and DNA integration (Table S1). Differential expression in the symbiont was also driven by the host developmental stage, with 77% (7,169) of all differentially expressed genes (9,355) regulated based on life stage (Fig S1b). Symbionts in adult corals overexpressed cellular metabolic processes, including photosynthesis, whereas symbionts in recruits overexpressed genes involved in reproduction, cell cycle processes, and DNA metabolic processes (Table S2).

**Figure 1:**
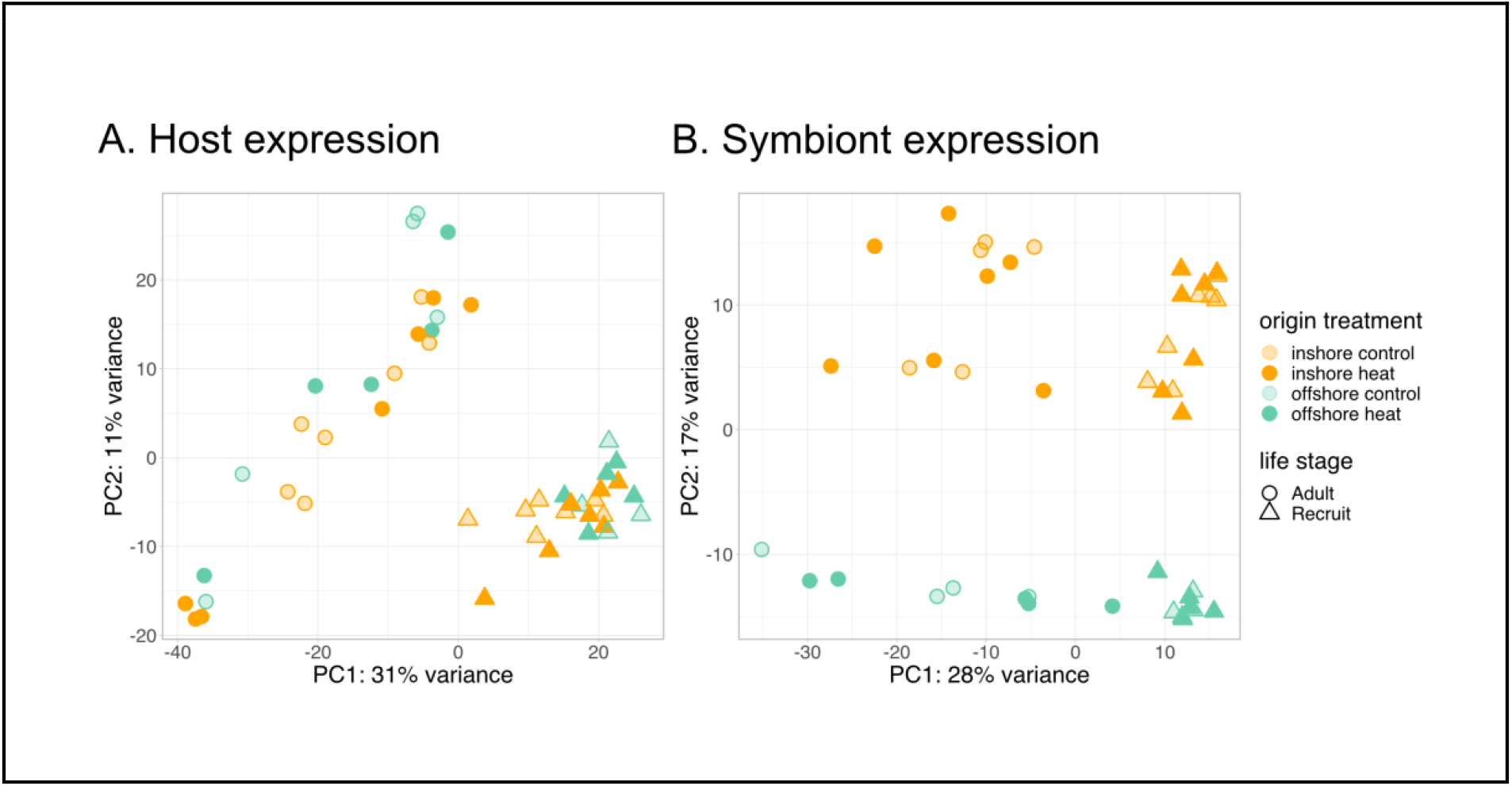
Principal components analysis of variance stabilized count data for the host (A) and symbiont (B) transcriptomes. Samples are colored by population and treatment. Symbols represent different life history stages.

### Signatures of reef origin are maintained, even in naive juveniles

Host and symbiont expression was also distinguished by reef origin regardless of life stage or treatment conditions. All host samples separated along the second principal component by origin, which explained ∼12.5% of the variance across life stages (Fig S2), despite earlier developmental stages never experiencing their home environments independent of the maternal colony. Reef origin accounted for the second highest proportion of differentially expressed genes across adult and recruit samples (1,302 genes, 15% of all DEGs; Fig S1a). Larval samples differentially expressed 225 genes due to origin, although treatment had a marginally larger effect (283 genes; Fig S3). Offshore corals exhibited greater expression of genes involved in carbohydrate transport, the TCA cycle, and nervous system development compared to inshore corals and reduced expression of genes involved in cellular component organization, localization, and vesicle-mediated transport (Table S3). Reef origin was also the primary driver of symbiont expression profiles along PC2 (Fig 1B; 17% of variance) and accounted for 1,700 (18%) of differentially expressed genes. No gene ontology terms were significantly enriched for origin-associated expression in symbionts.

### Moderate duration thermal stress induces moderate transcriptional regulation

Treatment accounted for the smallest number of differentially expressed genes in adults and recruits for both the host (605 genes, 7% of all DEGs; Fig S1a) and the symbiont (466 genes, 5% of all DEGs; Fig S1b). No discernable pattern was observed due to treatment across the first two principal components in either partner (Fig 1). A significant interaction was detected for 9 genes in which the treatment specific response differed according to the life stage of the host. Five of these genes were upregulated in adults and downregulated in recruits of which 2 were annotated: mitogen activated protein kinase 6 (MAPK6) and endothelin converting enzyme 2 (ECE2). In contrast, three of these genes were downregulated in adults and upregulated in recruits. One unannotated gene was uniquely upregulated in both life stages, but was more strongly regulated in adults compared to recruits (LFC 6.3 vs 1.4). No differentially expressed genes were detected for the origin by treatment interaction in the host, nor for any interaction terms in the symbiont. Although the majority of differentially expressed genes were regulated by treatment in larvae (283 genes, 56.5% of all DEGs), larval samples did not align along the first or second principal component by treatment (Fig S2). No genes were significantly differentially expressed due to the interaction of treatment based on larval origin.

### Treatment response is modulated by life stage

Although few genes were detected for the interaction of treatment and life stage, within group analysis revealed that the most treatment responsive genes were largely unique to each life stage. Recruits mounted the largest detectable expression response (311 DEGs) followed by larvae (283 DEGs) and then adults (194 DEGs). Only 9 genes consistently responded to treatment across all life stages (Fig 2C), including upregulation of the universal stress protein Sll1388, calumenin A, soma ferritin, and malate dehydrogenase. Conversely, expression of cyp1A1 varied, with adults and recruits upregulating cyp1A1 and larvae downregulating cyp1a1 under heat stress. The treatment response in symbionts was also modulated by the life stage of the host (Fig 3a). Symbionts in recruits had a greater expression response to treatment compared to symbionts in adults (155 DEGs versus 92 DEGs respectively) with only 16 DEGs overlapping between the two life stages (Fig 3b). These 16 core genes included annotations for fatty acid metabolism, RNA processing, metal ion binding, and oxidoreductase activity.

**Figure 2:**
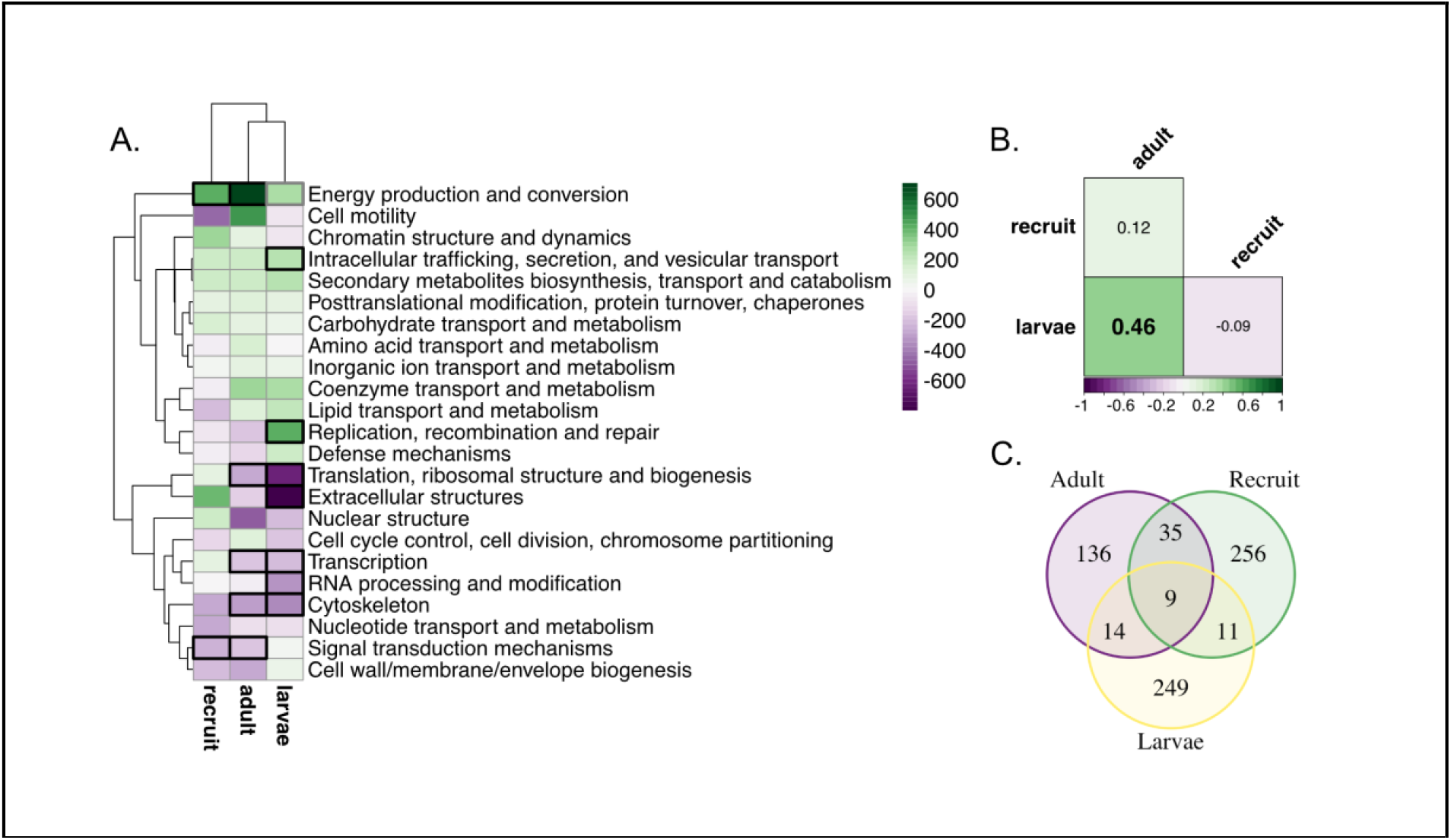
Host transcriptional response to moderate-term thermal stress by life stage. (A) Clustering of KOG enrichments by delta ranks and life stage. Bolded boxes represent KOG classes significantly enriched in response to treatment (black p_adj_ < 0.05; gray p_adj_ < 0.1). (B) Pairwise Pearson correlation coefficients of KOG delta ranks between life stages. Bolding denotes significant correlation (p < 0.05). (C) Venn diagram of genes significantly differentially expressed in response to treatment (p_adj_ < 0.1) in each life stage.

**Figure 3:**
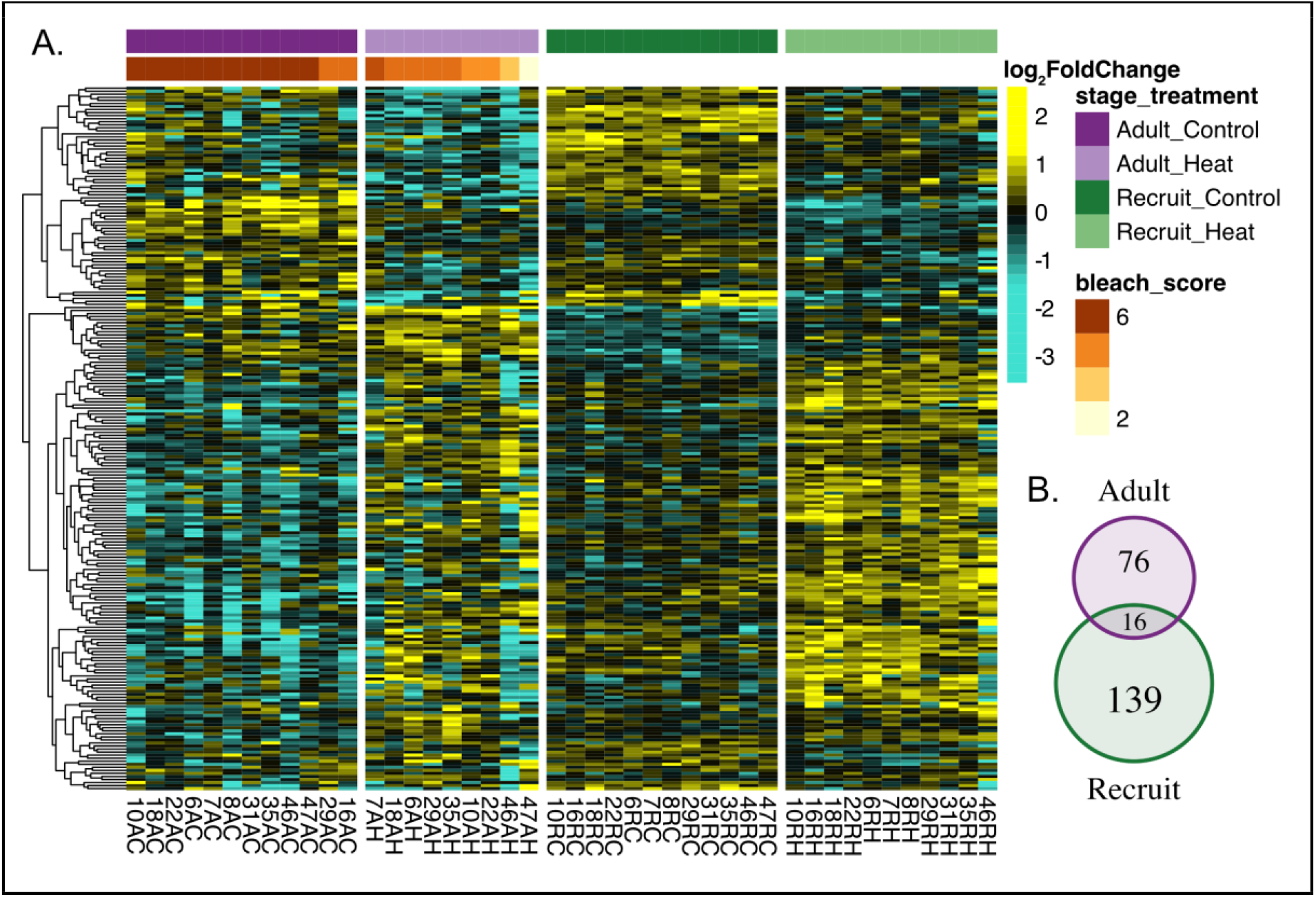
Heatmap of log_2_ fold change in expression of symbiont genes significantly differentially regulated in response to treatment across host developmental stages. Rows represent the 231 treatment responsive genes (Padj<0.1) detected in total for symbionts in adult and recruit life stages, as indicated by the venn diagram. Adult corals were ranked by visual bleaching score from 1 (lightest) – 6 (darkest) (Kenkel et al., 2015) which is denoted by the red/orange bar.

The unique differential expression response to treatment across life stages does not appear to be due to differences in variance (detection limits) but rather to differences in transcriptional baselines. Forty-four genes were constitutively expressed in adults, but responsive in recruits (Table S4). Whereas 27 treatment responsive genes that were unique to the adult life stage were overexpressed in control conditions by the recruit life stage (Table S4). Adults and recruits also exhibited backloading of 54 and 29 genes respectively. Symbionts exhibited a similar pattern of differing baselines leading to divergent thermal stress responses. Symbionts in adults frontloaded 15 genes and backloaded 28 genes that were responsive in recruit symbionts, while recruit symbionts frontloaded 22 genes and backloaded 17 genes that were responsive in adult symbionts (Table S5).

Functional enrichment analysis also indicates that life stages utilize different biological processes to respond to heat stress. Ontology enrichments in adults were driven by genes involved in upregulation of metabolic processes and downregulation of cellular amide metabolic processes (Fig S4a; Table S6), while recruits mainly downregulated genes involved in cell communication, membrane lipid biosynthetic processes, and cellular protein metabolic processes (Fig S4b; Table S7). Both adults and recruits upregulated organic acid catabolic processes and downregulated heterochromatin assembly by small RNA (Fig S4ab). Like adults, larvae also downregulated cellular amide metabolic processes. However, unlike either adults or recruits, larvae upregulated carbohydrate and DNA metabolic processes and downregulated cellular developmental processes (Fig S4c; Table S8). Among all life stages, energy production and conversion was the only KOG term significantly enriched among upregulated genes (Fig 2A). Although only 23 treatment responsive genes and 2 ontology term enrichments were shared among adults and larvae, a significant correlation in KOG class enrichment in response to treatment was evident (r = 0.46, p<0.05, Fig. 2B). However, KOG class enrichment was not significantly correlated in adults and their recruit offspring, in spite of high relatedness and exposure to identical treatment conditions (r = 0.12; Fig 2B).

Despite the lack of shared regulation at the level of individual genes, the biological processes generally upregulated by symbionts in response to heat stress were similar in each life stage, but downregulated processes varied. Both adult and recruit symbionts upregulated genes involved in RNA metabolism and cell cycle processes (Fig S5). Interestingly, recruit symbionts also upregulated genes involved in the cellular response to stress (Table S10), while adult symbionts downregulated photosynthetic processes (Fig S5; Table S9). Instead of photosynthetic processes, recruit symbionts downregulated nitrogen transport, utilization, and metabolism (Table S10).

### Treatment response is also modulated by reef origin, but patterns vary among life-stages

The magnitude of the expression response to heat stress also differed by population. Inshore adults and recruits had a much larger transcriptional response, differentially expressing 316 genes across treatments, whereas only 43 genes were detected as differentially expressed by offshore hosts (Fig 4A). Only 25 treatment responsive genes were differentially expressed in both populations. Universal stress protein 1388 (sll1388), Calumenin-A (calua), and malate dehydrogenase (MDH1) were responsive to treatment in both populations, as well as in all life stages, suggesting these may be core genes involved in the cellular stress response. Symbionts also exhibited population-level variation in the magnitude of their response to treatment, with inshore symbionts differentially expressing 142 genes compared to 57 genes in offshore symbionts, and only 14 genes showing consistent responses to heat stress between the two populations (Fig S6). Four of the five symbiont core genes annotated among populations were also identified as core genes among life stages: At2g29290, ppsD, PP1, and NUDT3. Regulatory protein flaEY, which plays a role in flagellum biogenesis, was also upregulated in response to heat stress in both symbiont populations.

**Figure 4:**
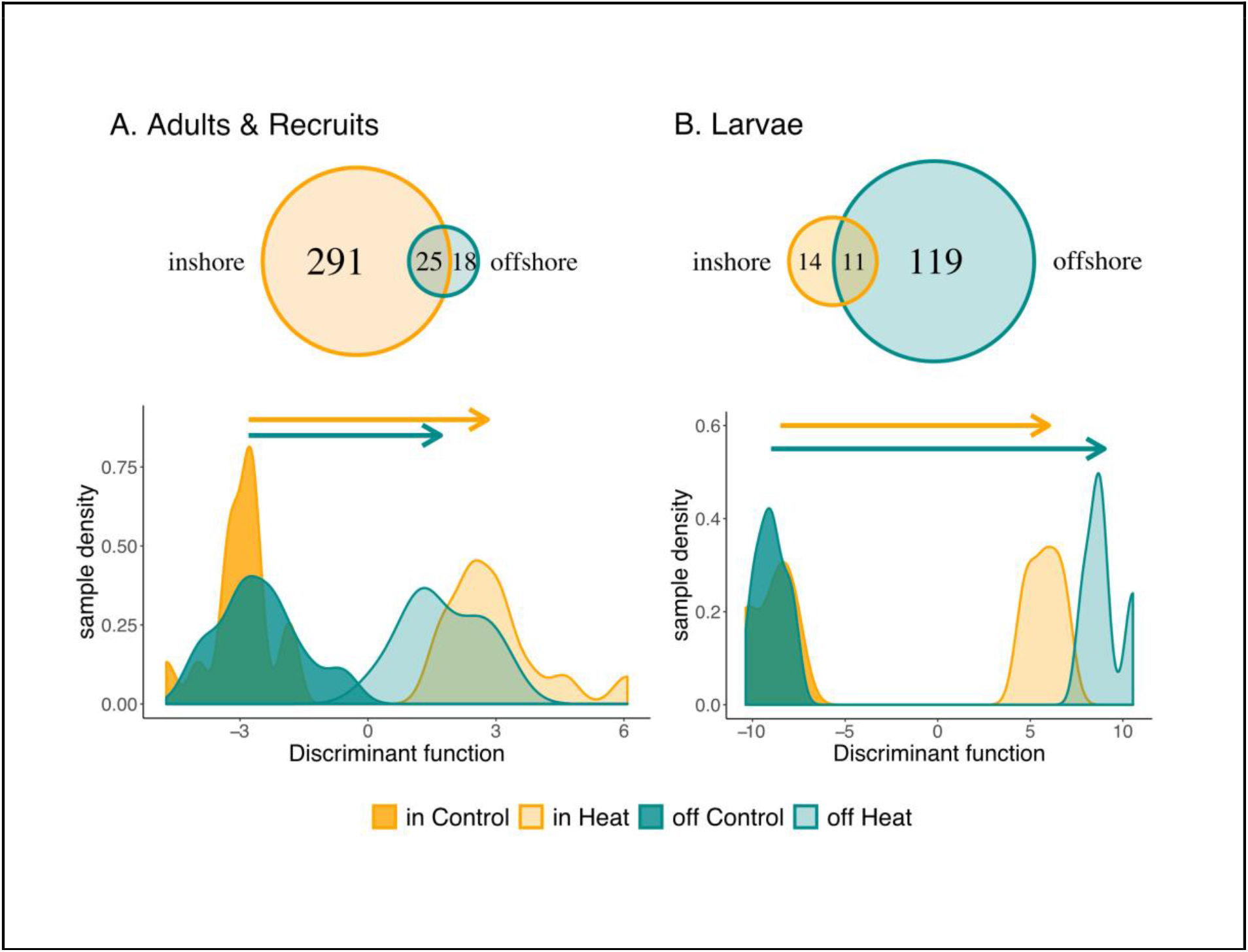
Host transcriptional response to treatment among populations. Venn diagram of the population-specific response to heat stress in adults and recruits (A) and larvae (B). Discriminant analysis of principal components was performed on all treatment responsive genes to explore the magnitude of expression change between populations. The discriminant function was defined by contrasting samples in heat and control from one population and predicting the response of the other.

Unlike the patterns observed for life-stage, the low number of treatment responsive genes in offshore adults and recruits does not seem to be due to front or backloading, but rather a reduced expression plasticity compared with inshore origin coral. Only 1 inshore responsive gene was frontloaded (LFC > 20 in control; Table S11) in offshore hosts, while 14 genes were backloaded (LFC −0.3 to −4.7 in control; Table S11). Offshore symbionts may be transcriptionally loading some inshore responsive genes (Table S12), with 11 genes strongly frontloading (fold change > 2) and one weakly backloaded (fold change < 2). However, for both host and symbiont, the total number of front and back-loaded genes in offshore samples is not enough to make up for the greater degree of differential expression in inshore samples.

Discriminant analysis of principal components for all treatment responsive genes supports a significantly muted expression response in offshore adults and recruits (P_MCMC_ < 0.01; Fig 4A). The robust inshore response in coral hosts included upregulated genes involved in catabolic and cellular metabolism and downregulated genes involved in actin-filament based processes and heterochromatin assembly. While DNA metabolic processes were upregulated in inshore corals, miRNA and protein metabolic processes were downregulated under heat stress (Table S13). Specifically, protein autoubiquitination was enriched among downregulated genes. Conversely, offshore hosts upregulated two stress response genes (ERP29 and slr1101) and one apoptosis gene (Mzb1). Although offshore-origin adults bleach significantly more than inshore-origin adults (Kenkel et al., 2015), the distance between adult samples along the discriminant function axis was not correlated with changes in bleaching score (Fig S7a).

Despite a consistent pattern of greater transcriptional response in inshore hosts and symbionts for adult and recruit life stages, larval hosts exhibited the opposite response. Offshore larvae differentially expressed 130 genes in response to treatment, whereas inshore larvae only differentially expressed 25 genes, with 11 genes shared among populations (Fig 4b). Offshore larvae also exhibited greater change in their transcriptional profiles for treatment responsive genes (DAPC, P_MCMC_<0.01; Fig 4b). Similar to adults, the distance between larval samples along the discriminant function was not correlated with changes in chlorophyll across treatments (Fig S7b). No genes were detected as front or back-loaded by inshore larvae.

## Discussion

Our results show that population origin and developmental stage have major impacts on gene expression and can modulate the response to heat stress in both coral hosts and their symbionts. A largely stage-specific thermal stress response implies inconsistent selection among life stages that could delay or confound adaptation. Despite differences among life stages, a fixed effect of population regardless of thermal history indicates a genetic basis for expression variation. Further, the magnitude of the treatment response varied based on population and was consistent across adults and their juvenile offspring, suggesting patterns of regulation are heritable. However, differences in response magnitude could also be influenced by the duration of heat stress or developmental stage necessitating further exploration. Considerable variation in baseline gene expression and the response to stressful conditions was also observed for symbionts, despite vertical transmission and similar ITS2 haplotypes between populations (Kenkel, Goodbody-Gringley, et al., 2013), implying that there is likely more physiological and/or genetic diversity in these communities than has been previously described.

### Developmental stage drives differential expression in coral hosts and modulates the thermal stress response

Life history stage was the prominent driver of transcriptional variation in the host (Fig 1a) and modulated the response to thermal stress (Fig 2c). Surprisingly, the thermal stress response was largely stage-specific, with few genes consistently responding to heat stress across adults, recruits, and larvae (Fig 2c). In contrast to the results of this study, Aguilar et al. (2019b) report a similar transcriptomic response to salinity stress between adult and juvenile corals. However, thousands of genes were exclusively responsive in a single life stage with only ∼20% DEGs consistently responding to salinity stress in this earlier study, reinforcing a largely stage-specific stress response (Aguilar et al., 2019b). Although sparse, there is also evidence for development modulating the stress response in plants (Garg et al., 2016; Liu et al., 2021) and other invertebrates (Devens et al., 2020; Schneweis et al., 2017), suggesting complex life cycles can decouple the molecular response to stress. The lack of consistency in the stress response across life stages has major implications for the evolutionary trajectory of these corals. If inducible genes in one life stage affect fitness, but are not expressed in other stages or are co-opted to perform another function, it may take longer for adaptation to occur, prolonging the evolutionary rescue which is predicted to play a key role in the persistence of corals in a rapidly changing environment (Bay et al., 2017b).

Stage-specific responses to thermal stress are partly due to differing baselines. Life history stage was the prominent driver of differential expression in the host (Fig S1), suggesting that genome-wide expression varies more over an individual’s lifetime than between locally adapted populations or different thermal environments. Developmental modulation of baseline gene expression could also influence stress responsive genes, potentially reducing plasticity during environmental perturbations. About one-third of treatment responsive genes in adults were front/back-loaded in recruits and one-third of treatment responsive genes in recruits were conversely front/back-loaded in adults. Front/back-loading of stress responsive genes has been proposed as an adaptive mechanism that increases coral thermal tolerance (Barshis et al., 2013b). However, if front/back-loading had a heritable genetic basis, offspring would be expected to exhibit constitutive expression similar to adults. Alternatively, if front/back-loading was environmentally-induced, it seems unlikely that thermally naive offspring would exhibit these patterns of regulation. Although it is possible that constitutive expression induces a priming effect against heat stress (Barshis et al., 2013b), the robust signal of developmental stage, regardless of genetic or environmental factors, more likely indicates that these genes are pleiotropic and perform different functions over developmental time.

Pleiotropic genes could have a positive effect on fitness in every stage of development, and thus increase total fitness, or they could experience divergent selection pressures over time, limiting adaptive potential (Cheverud et al., 1983). Pleiotropy is generally believed to constrain adaptation, however, a meta-analysis on quantitative genetic studies found genetic covariances were just as likely to promote, constrain, or have no effect on the rate of adaptation, although covariances among life stages were not considered (Agrawal & Stinchcombe, 2009). In model organisms, there is evidence suggesting that pleiotropy can constrain evolution through a change in the direction of selection across life stages. Quantitative trait loci in *Arabidopsis thaliana* can be positively correlated with fitness in one life stage, but negatively correlated in another (Postma & Ågren, 2016). Additionally, genes with stage-specific expression in *Drosophila* exhibit greater adaptive evolution than those expressed in multiple stages (Coronado-Zamora et al., 2019b). Although functional genomic evidence for pleiotropy is currently lacking for corals, bleaching resistance in adults can come at the cost of other fitness-related traits (Cornwell et al., 2021; Shore-Maggio et al., 2018), suggesting pleiotropy can constrain thermal adaptation within life stages. Putatively pleiotropic genes identified in the present study include tumor necrosis factor receptor superfamily 19 and a peroxidasin homolog, which have been previously implicated as key players in both the immune (Traylor-Knowles et al., 2021) and environmental stress response (Barshis et al., 2013b; Louis et al., 2017) in coral, yet their role in developmental modulation is unknown. The present study provides the first evidence for developmental pleiotropy in corals at a molecular level and emphasizes the need for functional and quantitative genetics to be conducted across multiple life stages to determine the costs and limits of thermal adaptation.

In contrast to one gene having multiple functions, different genes could also perform the same function, but be uniquely expressed during discrete developmental stages, resulting in a stage-specific thermal response. Transcriptome sequencing of five life history stages in two coral species, *Acropora palmata* and *Montastrea cavernosa*, found that expressed sequence tags (ESTs) were stage-specific (Schwarz et al., 2008), suggesting that life stages utilize completely different gene sets to achieve the same functions. Similarly, in *P. astreoides*, the inclusion of sequence data from three life-stages, larvae, newly settled recruits, and adults, increased the size of the transcriptome by 30 Mbp compared to the previously published adult-only reference (Mansour et al., 2016). As developmental stage was the main factor differentiating genome-wide expression in this study (Fig 1a), unique genes may be expressed during each life stage and result in differences in the identity of treatment responsive genes. Therefore, in addition to differing baselines, ontogenetic decoupling could also explain the lack of consistency in the thermal stress response among developmental stages.

Life stages could also be decoupled physiologically, rather than at the molecular-level, leading to potential differences in the susceptibility to thermal stress, and differences in the identity of treatment responsive genes. For example, adults may be more resilient to heat stress and be in an acclimatory phase, juveniles may be trying to reach homeostasis, and larvae could still be mounting a stress response due to the shorter duration of their exposure (Rivera et al., 2021). However, despite different genes, the response to heat stress in adults and larvae was functionally similar (Fig 2b), suggesting they may be in similar physiological states and deal with thermal stress in similar ways. The functional response of juvenile recruits, on the other hand, was not correlated with that of adults which experienced identical treatment conditions and are genetically related, suggesting the recruit life stage has a fundamentally different response from adults and larvae, and may experience stress differently. In support of this, Dixon et al. (2015a) also found a significant correlation in the rank of KOG classes enriched in *A. millepora* adults and larvae in response to elevated temperatures, although recruit responses were not explored. Physiologically, recruits may be investing more energy in individual growth, while adults use this energy for sexual reproduction. However, differentially expressed genes between these two stages were not enriched for reproductive processes (Table S1), indicating that there are likely other physiological and functional differences that vary across developmental stages. Future work needs to focus on bleaching phenotypes that can be compared among all stages to establish the resilience/susceptibility of different developmental stages to controlled thermal stress.

Ontogenetic decoupling could delay adaptation because expression of specific genes will only have fitness effects during discrete developmental periods (Albecker et al., 2021). However, decoupling could also break genetic correlations between discrete life stages, reducing antagonistic pleiotropy, and therefore optimizing stage-specific fitness (Moran, 1994). As metamorphosis is often accompanied by changes in habitat, ontogenetic decoupling is predicted to be strongest across metamorphic transitions (Moran, 1994). Interestingly, in addition to differences in the thermal stress response between larvae and post-metamorphic life stages, we also see clear differences in the identity of treatment responsive genes between recruits and adults (Fig 2c), which lack any major morphological or ecological transitions. This suggests that adults and recruits, in addition to larvae, vary in their fitness optima and can evolve somewhat independently. In *Drosophila*, artificial selection on larval or adult tolerance to extreme temperatures did not confer tolerance in the other life stage (Loeschcke & Krebs, 1996) and the genetic architecture underlying tolerance in each stage was found to be independent of the other (Freda et al., 2017; Loeschcke & Krebs, 1996). A largely decoupled thermal stress response among all three life stages in this study suggests that selection for tolerant coral genotypes in one life stage will likely not translate into greater tolerance in subsequent stages, as the genes under selection will not be expressed. Human intervention strategies, such as artificial selection and assisted gene-flow, have been proposed to save vulnerable coral populations from bleaching events (van Oppen et al., 2015), but our results suggest further research is needed to determine the effect of genetic variation on total organismal fitness, including all life stages, before these can be successfully implemented.

### Population-level signatures persist in thermally naive life stages

Origin effects on constitutive expression persisted in all life stages despite rearing adults in a common garden for five weeks and larvae and recruits having no exposure to their home environment independent from the maternal colony. Although adult *P. astreoides* from inshore and offshore reefs in the Florida Keys were previously found to exhibit population-level divergence (Kenkel, Goodbody-Gringley, et al., 2013; Kenkel, Meyer, et al., 2013; Kenkel & Matz, 2016), the underlying mechanisms driving expression variation could not be determined. Here we see clear differentiation between inshore and offshore expression profiles in juvenile life stages reared in a common environment, suggesting a genetic basis for expression divergence. Population-level differences were also present in the thermal stress response, where inshore adults and their recruit offspring exhibited a larger shift in expression profiles for all treatment responsive genes and greater differential expression across treatments than offshore-origin adults and recruits (Fig 4a). Inshore adults of *P. astreoides* also exhibit greater genome-wide expression plasticity in response to transplantation (Kenkel & Matz, 2016) and are generally more thermally tolerant compared to offshore adults (Kenkel et al., 2015; Kenkel, Goodbody-Gringley, et al., 2013). Here we show that this expression pattern is consistent in juvenile offspring, with no prior thermal stress experience, indicating that plasticity is potentially heritable.

However, origin specific differences in expression could also be due to transgenerational effects induced by the parental environment, including maternal effects or developmental plasticity. Some studies have documented an effect of parental environment on offspring physiology in coral (Bellworthy et al., 2019; Putnam & Gates, 2015; Wong et al., 2021), suggesting that underlying expression patterns likely play a role. It is important to note that because *P. astreoides* employ a brooding reproductive strategy, larvae and juveniles in this study were exposed to their home environment during embryonic development. Temperature profiles are similar between these inshore and offshore reefs in the fall and spring when embryonic development occurs, yet many other environmental parameters diverge (Kenkel et al., 2015), which could have induced a plastic phenotypic response. Though we cannot rule out maternal effects or developmental plasticity, brooded offspring are commonly retained within reefs (Ayre & Hughes, 2000; Underwood et al., 2007), so persistent population-level differences could facilitate survival of early life stages in their local environment.

Despite population-specific differences being consistent in parents and offspring, expression plasticity may not be the primary mechanism driving differences in thermal tolerance. Similar to adults, inshore larvae are more resistant to bleaching than offshore larvae (Zhang et al., 2019). However, opposite to the population-specific plasticity observed in adults and recruits, larvae from offshore families demonstrated a larger shift in expression profiles for all treatment responsive genes and differentially expressed more genes across treatments compared to inshore-origin larvae (Fig 4b). Reduced expression plasticity due to constitutive expression of thermally responsive genes, or front/back-loading, has been documented in tolerant coral adults in American-Samoa (Barshis et al., 2013b) and the Red Sea (Voolstra et al., 2021). However, we did not detect any front/back-loaded genes in inshore larvae, suggesting this is not the main mechanism driving population variation. Additionally, expression plasticity was not correlated with plasticity in adult bleaching score (Fig S7a) or larval chlorophyll content (Fig S7b), implying that population-level differences in plasticity do not predict the magnitude of bleaching.

Alternatively, the timing of the cellular stress response may vary across populations and underlie differences in thermal tolerance rather than its plasticity (Rivera et al., 2021). Population-specific plasticity in larvae exposed to only 4 days of heat stress was opposite that of adults and recruits exposed to 16 days of heat stress. As offshore-origin larvae had a more robust response to acute stress (Fig 4b), but inshore-origin adults and recruits had a more robust response to moderate heat stress (Fig 4a), the onset of the stress response might occur earlier in offshore corals, artificially leading to differences in population-specific plasticity across life stages. It is also possible that after 16-days of moderate thermal stress, population-specific responses observed in adults and recruits are due to differences in cellular fate, rather than the cellular stress response. Treatment responsive genes in inshore adults and recruits were mainly involved in metabolism, actin-filament based processes, and heterochromatin organization (Table S13), whereas the offshore-specific response included genes involved in the cellular stress response and apoptosis. This pattern could indicate that offshore adults and recruits may have reached their thermal limit and activated cellular death pathways, while inshore corals are utilizing other pathways to acclimate to warmer conditions. Multiple sampling timepoints during controlled thermal stress are needed to resolve whether differences in response timing or plasticity underlie differences in cell-fate and affect the bleaching response.

### Symbiont expression is modulated by host and holobiont condition

Symbiont expression patterns reflected those exhibited by the host, with life stage and population of origin both differentiating expression and modulating the response to heat stress. Stage was the prominent driver of transcriptional variation (Fig 1b), providing the first evidence that symbiont expression is affected by the host’s developmental stage. Symbiont community composition is presumed to be similar across life stages and populations, as symbionts are directly inherited by recruits through the process of vertical transmission and the same symbiont haplotypes (ITS2 type A4/A4a) were previously found in adult *P. astreoides* at these reef sites (Kenkel, Goodbody-Gringley, et al., 2013), consistent with broader geographic patterns (Thornhill et al., 2006). Consequently, high transcriptional variation among similar symbiont communities could be caused by differences in the host environment. However, symbiont identity can also drive expression variation (Berkelmans & van Oppen, 2006; Rowan, 2004; Rowan et al., 1997; Sampayo et al., 2008), so it is possible that there is more cryptic genetic variation in symbionts than has previously been described. Interestingly, constitutive differences in symbiont expression also led to a largely stage- and population-specific response to thermal stress, suggesting the susceptibility to heat stress could vary based on host-symbiont interactions.

If symbiont populations are genetically homogenous, expression variation could be due to changes in the relationship with hosts over development. In newly released planula larvae, symbionts translocate comparatively less carbon to their hosts than adults (Kopp et al., 2016), but up to 70% of photosynthetically fixed carbon is translocated by symbionts in 22-27 day old planulae, similar to adult symbioses (Gaither & Rowan, 2010). Because symbiont densities also increase as larvae mature (Kopp et al., 2016), symbionts in early life stages may invest more energy into population growth, but then translocate more carbon to hosts when symbiont population density stabilizes. Interestingly, functional enrichments identified here show that symbionts in recruits overexpressed genes involved in reproduction, DNA replication, and cell cycling compared to symbionts in adults, but underexpressed genes involved in photosynthesis (Table S2), implying that even after metamorphosis, symbiont populations in 7-week old recruits are still investing energy in population growth. This suggests that the metabolic balance between host and symbiont varies over development, even when symbionts are vertically transmitted, and could have implications for how symbiosis is maintained during different developmental stages.

Reduced baseline expression of photosynthetic genes in juvenile life stages could also reduce oxidative damage at high temperatures and increase bleaching resistance (Lesser, 1996). Under heat stress, symbionts in recruits upregulated cellular stress response genes while symbionts in adults downregulated genes involved in photosynthesis (Fig S5). Expression of photosystem II reaction center protein L (psbL) was negatively correlated with bleaching in adults (Fig S8), suggesting that downregulation of photosynthetic genes in heat-stressed adults is likely due to photoinhibition rather than an acclimatory response. Reduced expression of other PS II core proteins, such as psbA, have been previously detected in thermally sensitive Symbiodiniaceae species and are proposed to limit the repair rate of PSII under heat stress (Gierz et al., 2016; McGinley et al., 2012). In contrast, psbL was more moderately regulated in recruits (Fig S9), suggesting symbionts in recruits may be actively compensating for oxidative stress. Therefore, juvenile holobionts may be more resilient to thermal stress than their adult counterparts, but comparable bleaching phenotypes must be measured across life stages to test this hypothesis.

An alternative explanation for the life-stage specific expression could be that symbiont communities in juveniles differ from those in adults. A growing body of literature suggests that vertical transmission may not be as exclusive as previously thought. In three heritable symbioses, early life stages contained a higher diversity of background symbiont types that were not present in maternal colonies, indicating the potential for mixed-mode transmission (Quigley et al, 2017; Quigley et al, 2018; Byler et al, 2013). In our host expression dataset, recruits under-expressed immune-related genes compared to adults (Table S1), which could facilitate the uptake of exogenous symbionts. Evidence for fixed differences in symbiont expression has been reported at the genus, species, and strain-level (Barshis et al, 2014; Barfield et al, 2018; Parkinson et al., 2016), so differences in symbiont expression across life stages observed here could be due to changes in community composition across development. Although symbiont communities in adult populations of *P. astreoides* appear to be conserved (Kenkel, Goodbody-Gringley, et al., 2013), no study has yet to explore potential changes across generations, warranting further exploration in this species.

Strain-level variation could also diverge between reefs, leading to population-level expression variation in both adult and recruit life stages (Fig 1b). Although the major haplotypes hosted by adult *P. astreoides* have been previously reported to be similar across these sites (Kenkel, Goodbody-Gringley, et al., 2013), vertical transmission could cause local retention of symbiont strains within each population leading to genetic differentiation. After a year-long reciprocal transplant of *P. astreoides* adults across these same populations, non-native symbiont expression profiles shifted towards that of the native population, but were still distinct, indicating the existence of fixed differences between symbiont populations (Kenkel & Matz, 2016). Maximum and effective quantum yield of PSII of symbionts from these same populations in-hospite was also higher in inshore-origin adults under heat stress compared to offshore-origin adults (Kenkel, Goodbody-Gringley, et al., 2013), which could be explained by the more robust transcriptomic response to heat stress in inshore-origin symbionts observed here (Fig S6). Alternatively, symbiont expression could also differ due to long-term acclimatization to the host environment. However, symbionts in environmentally naive recruits also exhibit population-level differentiation, indicating genetics of the symbiont and/or host are more likely driving this variation. As host populations were previously found to be genetically differentiated (Kenkel, Goodbody-Gringley, et al., 2013), we cannot rule out the possibility that local adaptation of the host is driving differences in symbiont expression across reefs. Fine-scale population genetics is needed to test whether genetic differentiation in the symbiont is a possible mechanism for population-level expression divergence observed here.

## Acknowledgements

CDK was supported in part by fellowship funds from the Alfred P. Sloan Foundation.

