## Supplemental figures for "Divergent transcriptional response to thermal stress among life stages could constrain coral adaptation to climate change"

S1. Venn diagram of significant differentially expressed genes by life stage, reef origin, and treatment for (A) host expression and (B) symbiont expression in adult and recruit corals.

A.	B.

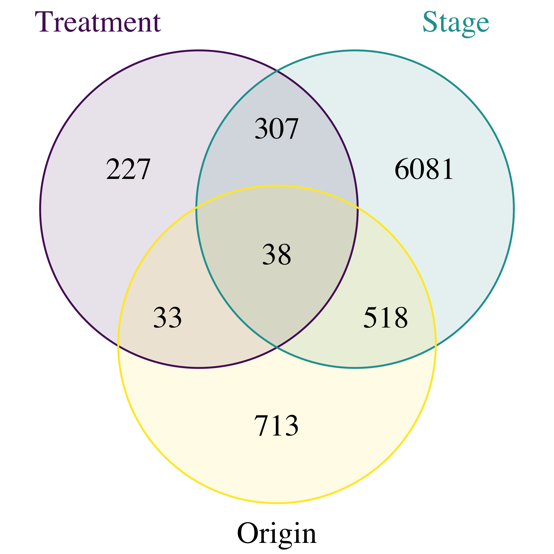

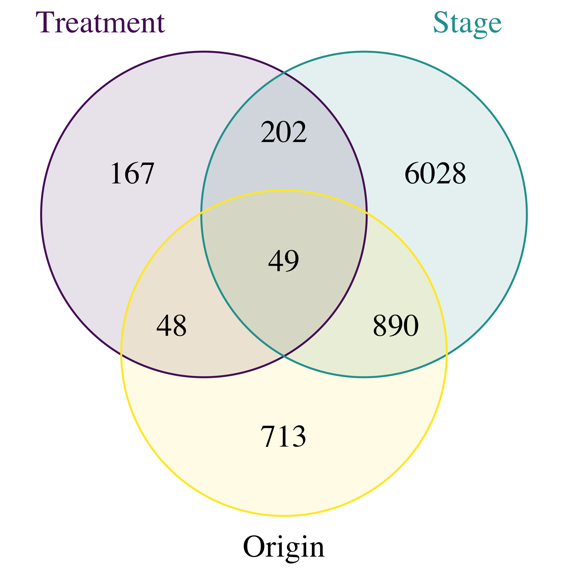

S2. Principal components analysis of adult, recruit, and larval host expression.

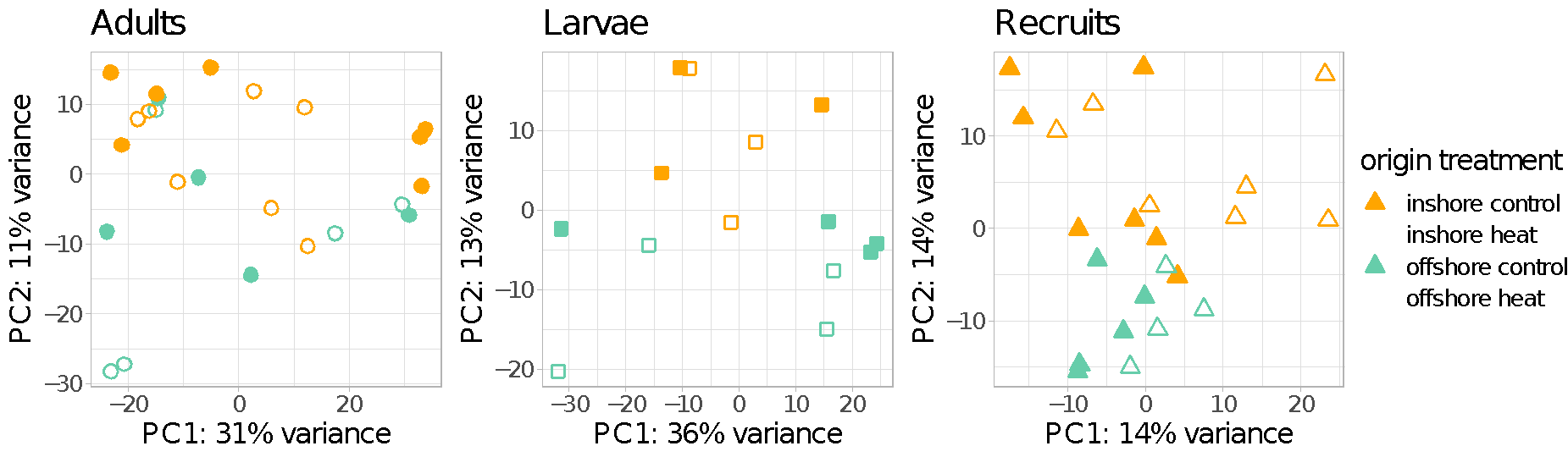

S3. Venn diagram of significant differentially expressed genes by reef origin, and treatment for host expression in larvae.

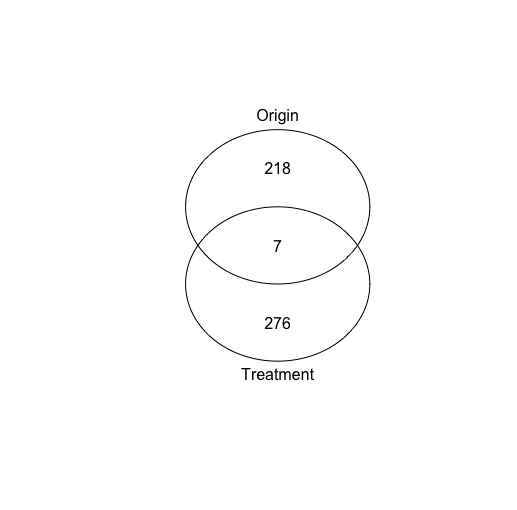

S4. Gene ontology enrichments for treatment responsive genes in each coral life stage: (A) adult, (B) recruit, (C) larvae.

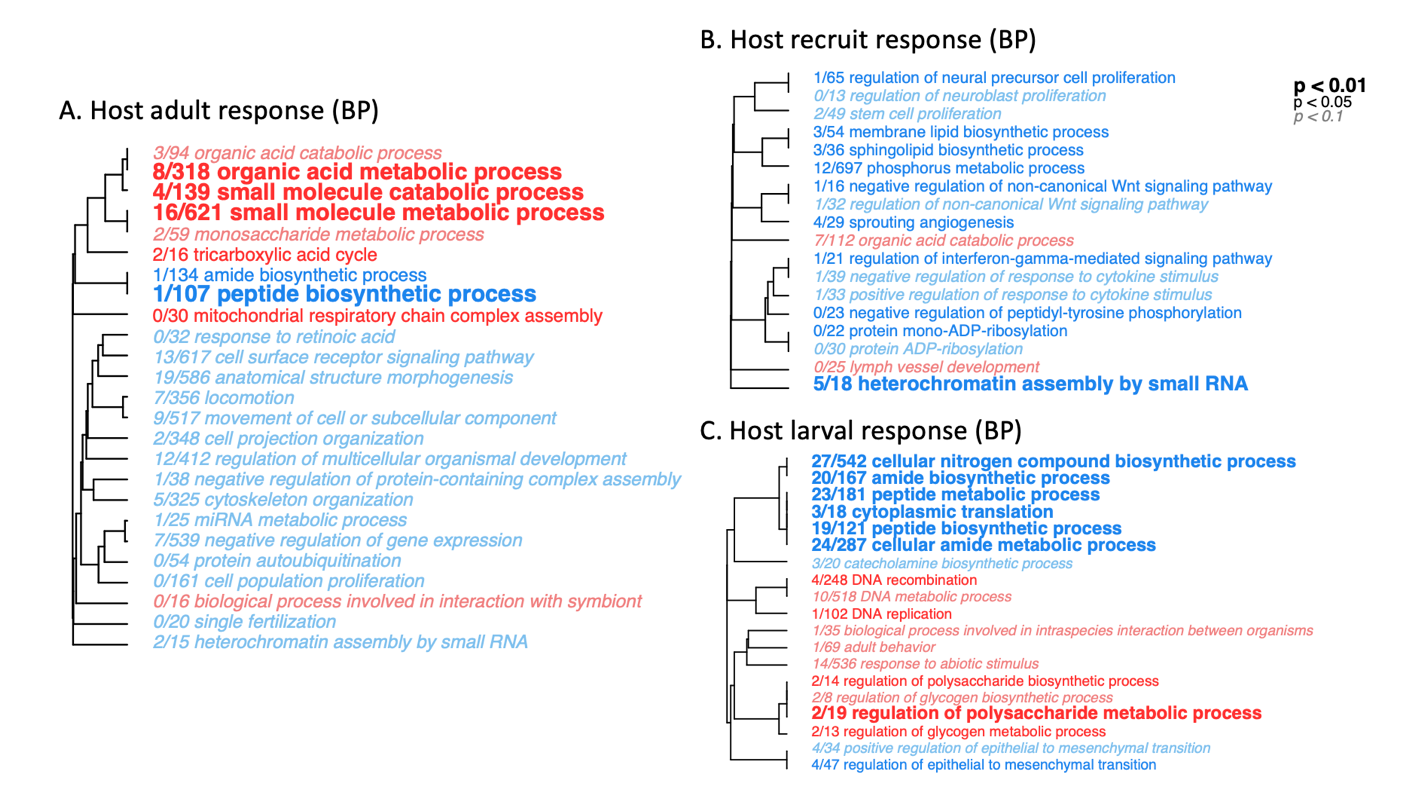

S5. Gene ontology enrichments for treatment responsive genes in symbionts inhabiting different life stages of its host: (A) adult and (B) recruit.
A. Adult symbiont response

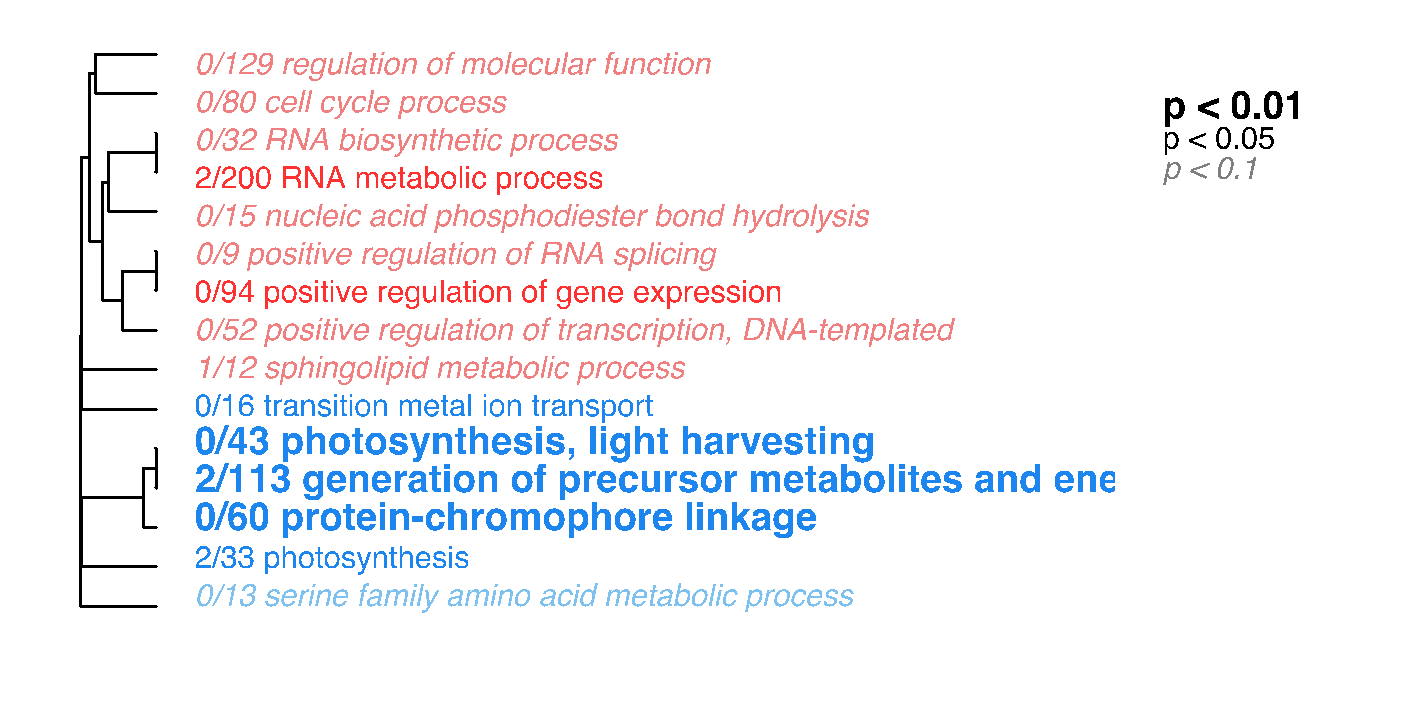

B. Recruit symbiont response

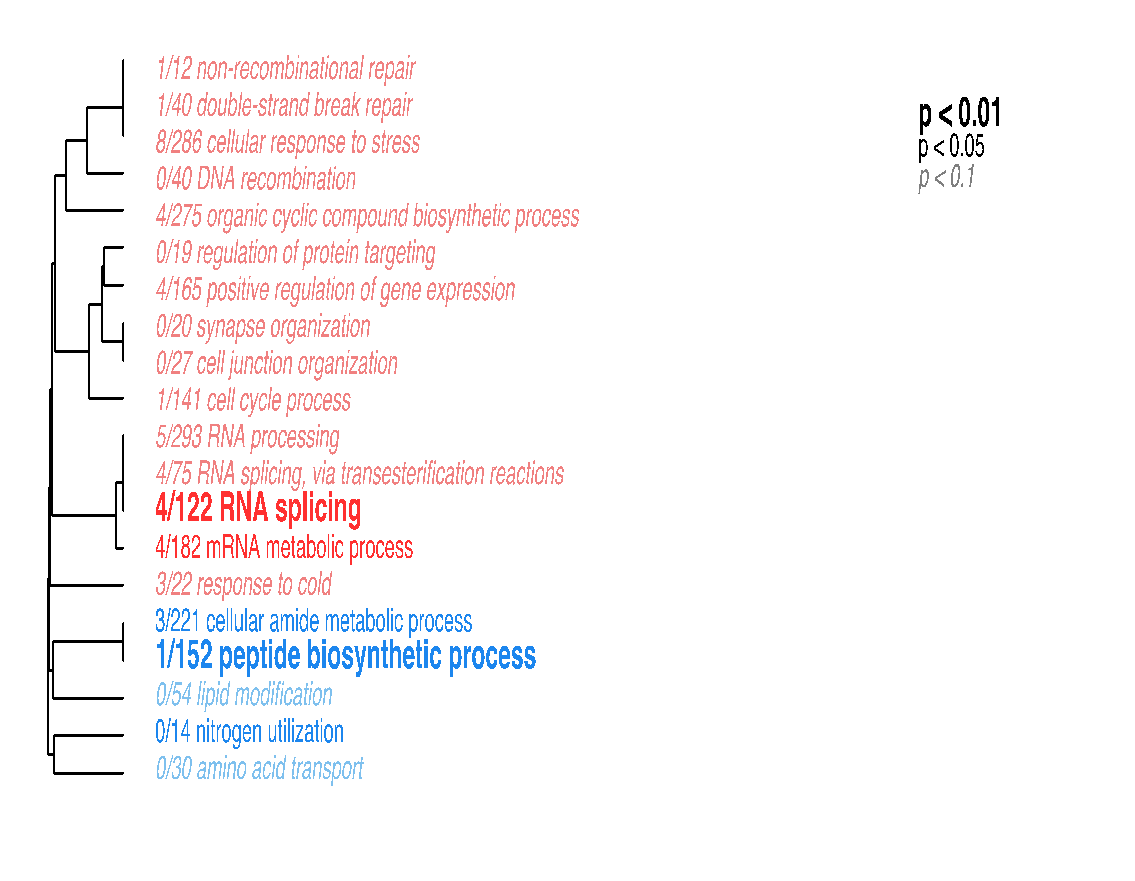

S6. Venn diagram of treatment responsive genes in the symbiont by population.
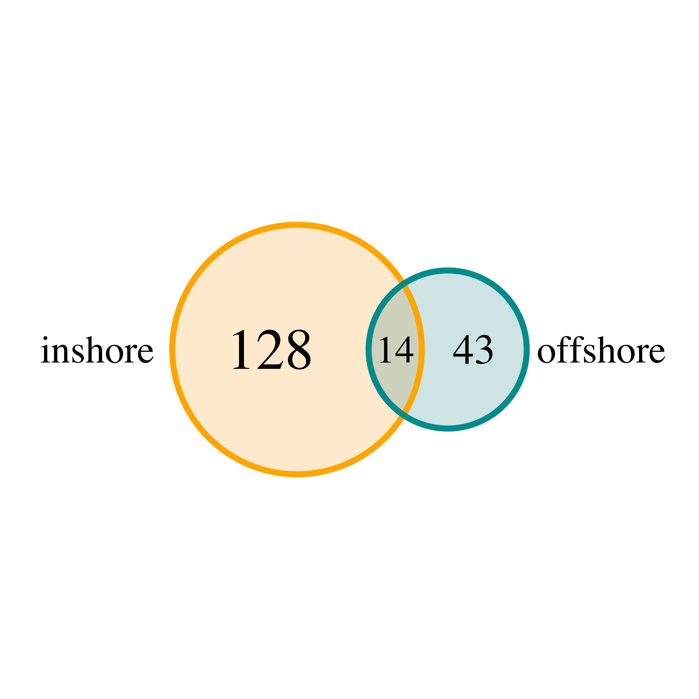

S7. Correlation of plasticity in thermal stress response to plasticity in bleaching score of adults (a) and larval chlorophyll content (b).

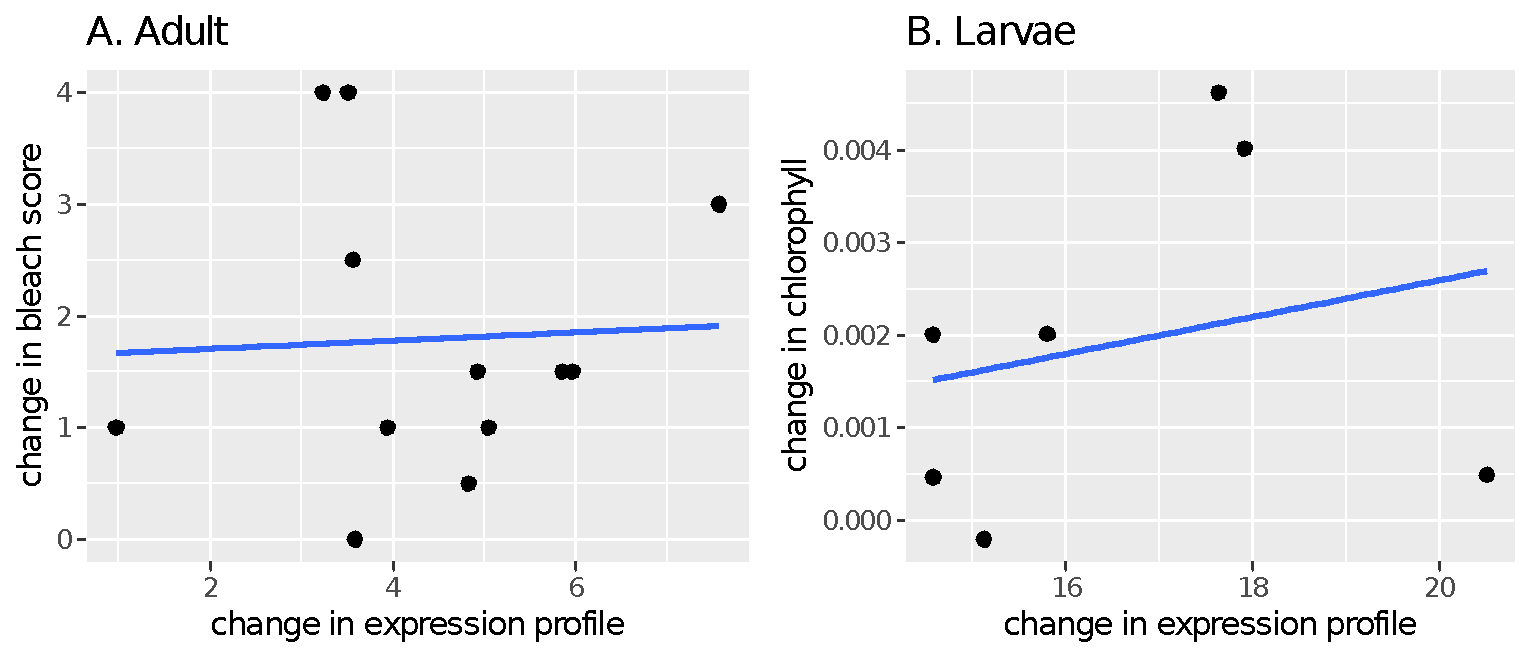

S8. Expression of photosystem II reaction center protein L (psbL) is correlated with bleaching score in adults.

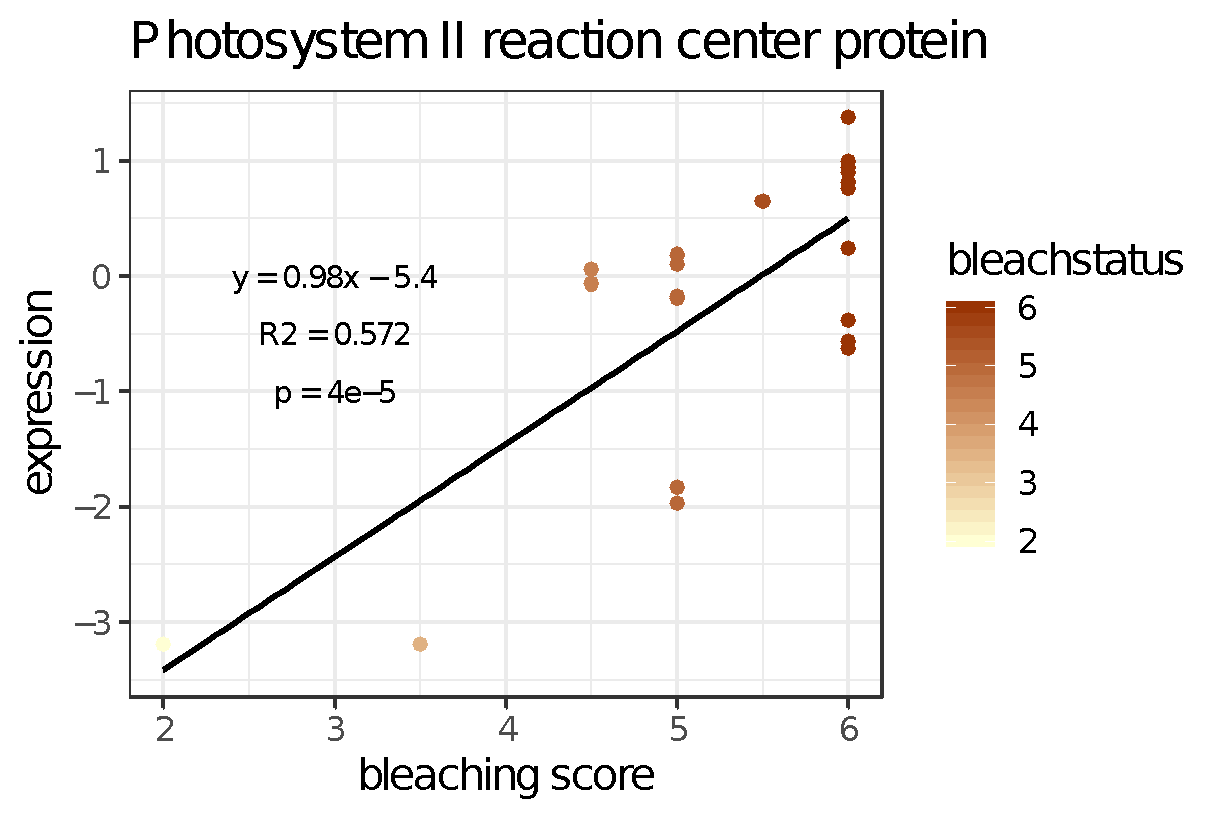

S9. Heatmap of photosystem II reaction center protein (psbL) expression of symbionts in adults and recruits exposed to heat stress.
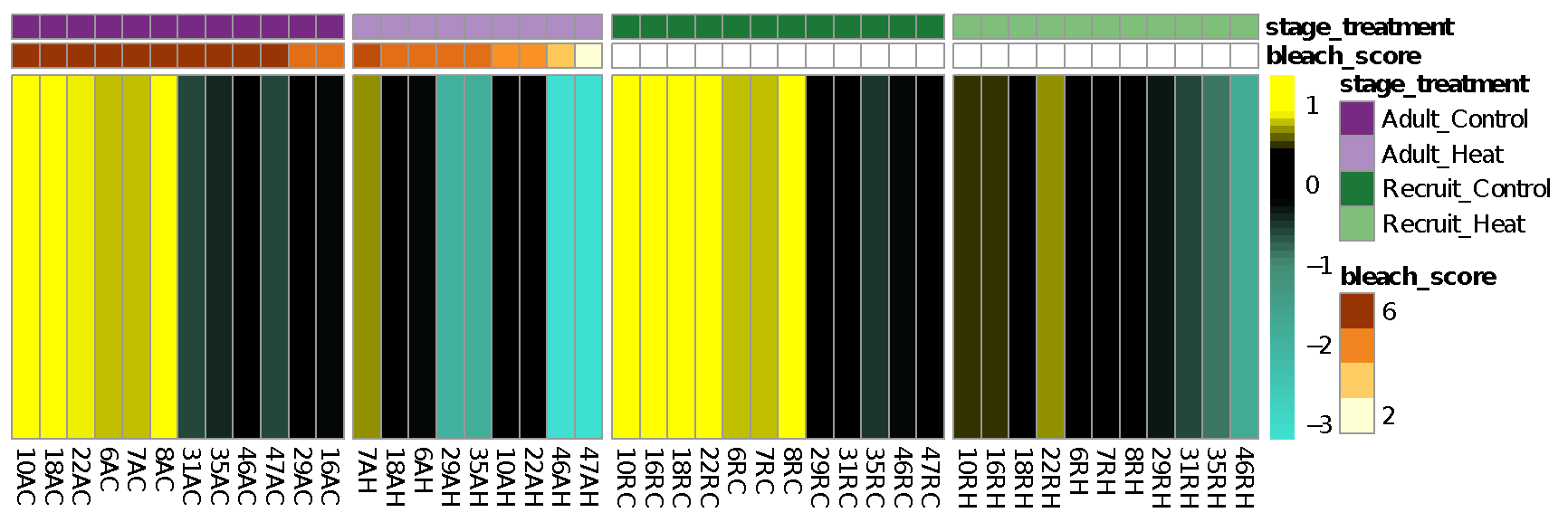
